# *Athene cunicularia hypugaea* wintering in a central California urban setting arrive later, leave earlier, prefer sheltered micro-habitat, tolerate rain, and contend with diverse predators

**DOI:** 10.1101/2024.09.17.613585

**Authors:** Martin A Nicolaus

## Abstract

The wintering phase of the Western Burrowing Owl (*Athene cunicularia hypugaea*) life cycle has received little attention in the literature. Small numbers of Burrowing Owls have been observed and recorded since 2008 in the winter season in Cesar Chavez Park, a 36-ha peninsula that forms part of the waterfront of the City of Berkeley. The relative ease of observation in this setting allows study of their arrival and departure dates, selection of micro-habitats, tolerance of human presence, behavior in inclement weather, and response to avian and canine predator threats. Viewed over a decade, fewer owls arrived, arrived later, left earlier, and spent less time in residence. Most owls settled in shoreline rip-rap or in tall vegetation; only one fourth settled in short grass. Owls chose exposure to rain for as long as two days. Owls varied widely in tolerance to human proximity. Owls successfully dealt with avian predators, but displayed stress and in some cases became casualties of loose *Canis lupus familiaris*. Recommendations for habitat management follow.

## INTRODUCTION

The Burrowing Owl (*Athene cunicularia hypugaea*) is a Species of Special Concern in California. (Wilkerson and Siegel 2010) A petition to classify it as endangered or threatened is pending before the California Fish and Game Commission. (Center for Biological Diversity et al. 2024)

Most research about Western Burrowing Owls (*Athene cunicularia hypugaea*) has focused on their breeding ecology. As early as 1990, Haug and Oliphant noted that “migration routes and wintering areas are unknown.” (Haug and Oliphant 1990) The same observation was made in subsequent years (James 1992; Klute et al. 2003; Lantz, Smith, and Keinath 2004; Woodin, Skoruppa, and Hickman 2007; Lincer et al. 2018). Despite some recent advances in winter ecology in Texas and in Mexico, Hall and Conway were able to write as recently as 2021 that “little is known about their winter and migration ecology including where they spend the winter, migration routes, and stopover sites.” (Hall and Conway 2021)

## METHODS

A small number of wintering Burrowing Owls have visited Cesar Chavez Park on the waterfront of the City of Berkeley since the mid-1980s. From 2010 through 2017, a docent program organized by the local Audubon chapter trained and dispatched volunteer observers to record owl sightings periodically, and these were summarized in annual reports. From 2018 on, a citizen-science project organized by the Chavez Park Conservancy mobilized park visitors to visit the owl habitat on a daily basis, reporting finds to the chavezpark.org website for publication, usually the same or next day. More than two dozen park visitors contributed reports and/or photographs. These efforts yielded more than 300 daily narrative reports with photographs published on the chavezpark.org website and more than 250 YouTube videos of Burrowing Owls in this location. Together, the docent documentation and the citizen-science reports provide the basis for the current study.

Observation in Chavez Park is in many ways easier than field observation in the wilderness. The park lies about fifteen minutes by car from the Berkeley city center. One could walk from the nearest parking area to the owl habitat in about 10 minutes, and cover the entire area where owls have been seen in about 15 minutes on foot. A few owls could be easily seen with the naked eye. Others required long-range optics. A number of owls settled in dense tall grass or behind screens of fennel, and observers spent considerable time systematically scanning the areas to spot owls concealed there.

This was a citizen-science effort. Invasive techniques such as trapping the birds, attaching bands or radio collars, digging out burrows, etc., were not used here, nor were audio call-broadcasts and devices such as Robel poles or Daubenmire frames.

## THE HABITAT

Cesar Chavez Park is a municipal park located on the waterfront of the City of Berkeley, California, in the western portion of Alameda County. (Fig. 1) The weather is generally Mediterranean and marked by persistent winds, with a prevailing west and northwest daytime influence. South, north, and east winds also occur. Mean temperatures in the fall and winter months (Oct - Mar) average between 13°C and 17°C. Snowfall is absent. Rainfall, when it occurs, is concentrated in the winter months (Nov - Mar), but varies greatly. The winter years 2016-2021 were marked by drought. The winters 2021-2022 and 2022-23 saw extreme rainfall. (NOAA 2024)

**Fig. 1.**
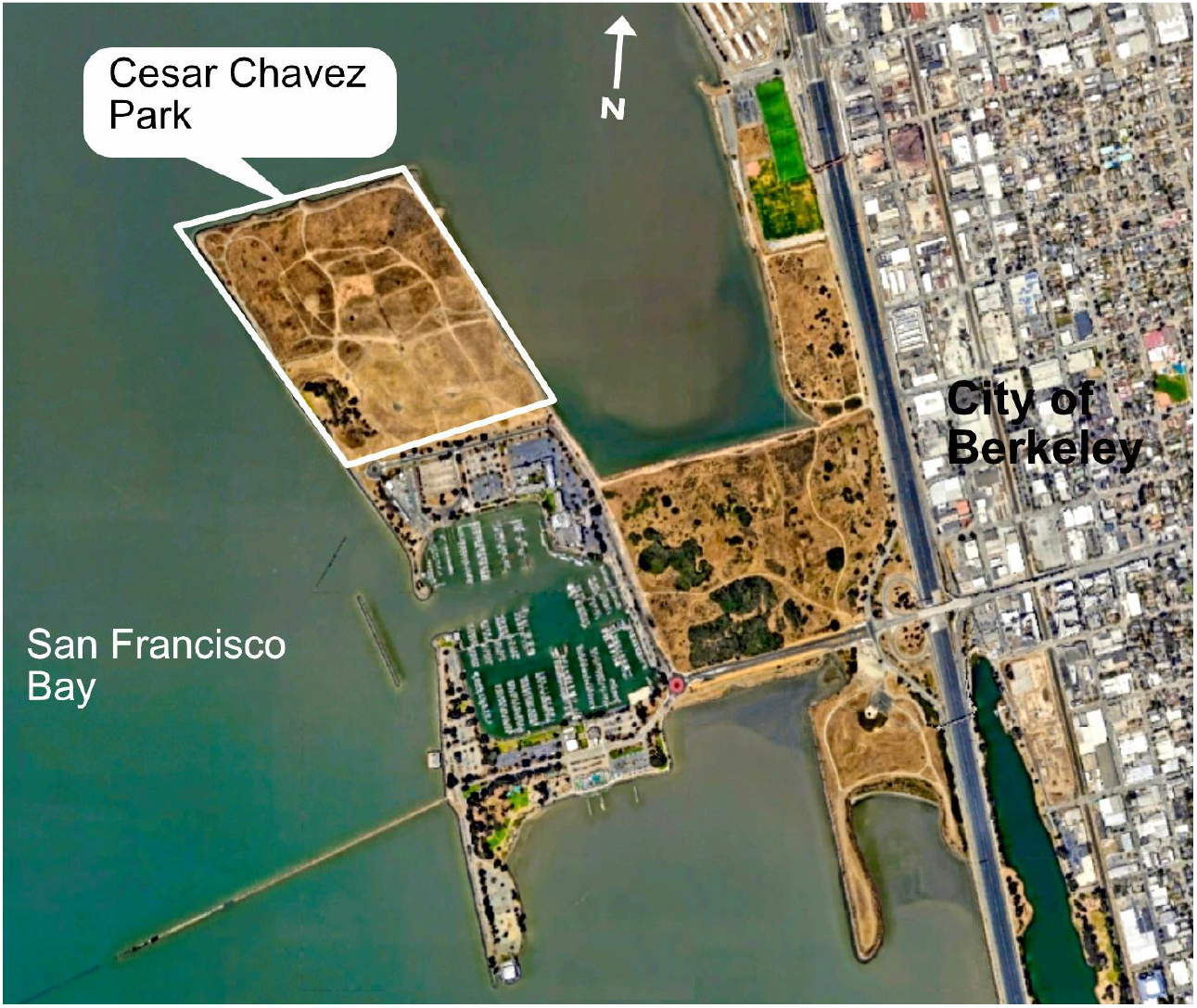

The park is a former landfill erected on tidal mudflats. Its west, north, and east borders consist of a human-made sloping embankment of stones (rip-rap) that raise the base of the park about 4m above the mud base. Beginning in 1961, the enclosure served as the city dump and was filled with municipal solid waste until 1983. It was then capped and converted into a park. (Minshew and Sison 2021) The surface area is about 36 ha. The south boundary is a two-lane paved road, Spinnaker Way, that fronts a 437-room hotel, some parking areas, and a boat maintenance and repair yard. On its east side, an estuary, the North Basin, about 650m wide on the average, separates the park from the city’s mainland. The north side faces a branch of San Francisco Bay, bounded by the port of the City of Richmond about 4.5km distant. The west side features bay views of the City of San Francisco (∼10km), the Golden Gate Bridge (∼15km), and the coast range to the north.

The terrain consists of gentle slopes rising from east to west, with a west ridge rising about 20m above the base. The north and northeast edges feature a flat grassland sloping softly to a southern ridge about 4m higher. A mostly level paved path about 2.3km long forms a loop along the entire perimeter for pedestrian and bicycle use and is the main visitor access route.

The area is exposed to fairly constant freeway noise from Interstate 580 about 1000m east, an iron forge plant about 1100m east, and railroad whistles approximately 1300m east. Passenger jets approaching San Francisco and Oakland airports can be heard regularly. The human park visitor quickly learns to tune these noises out and focus instead on the sounds of nature. The extent to which the noise level disturbs wildlife is an unexplored dimension in this setting.

The western portion of Alameda County showed some breeding populations of Burrowing Owls historically (Townsend and Lenihan 2003). But there is no indication that Burrowing Owls ever nested in the park or in its predecessor, the landfill. All the BUOW seen here were migrants, either as seasonal settlers or as stopovers. An eBird hotspot located in the southeast corner of the park is nominal and does not represent actual owl locations (Cornell Laboratory of Ornithology 2024). All the Burrowing Owls recorded in the park since 2008 settled at various points along the north shore.

Three defined areas along the north shore have served visiting owls as seasonal or stopover habitat. (Fig. 2)

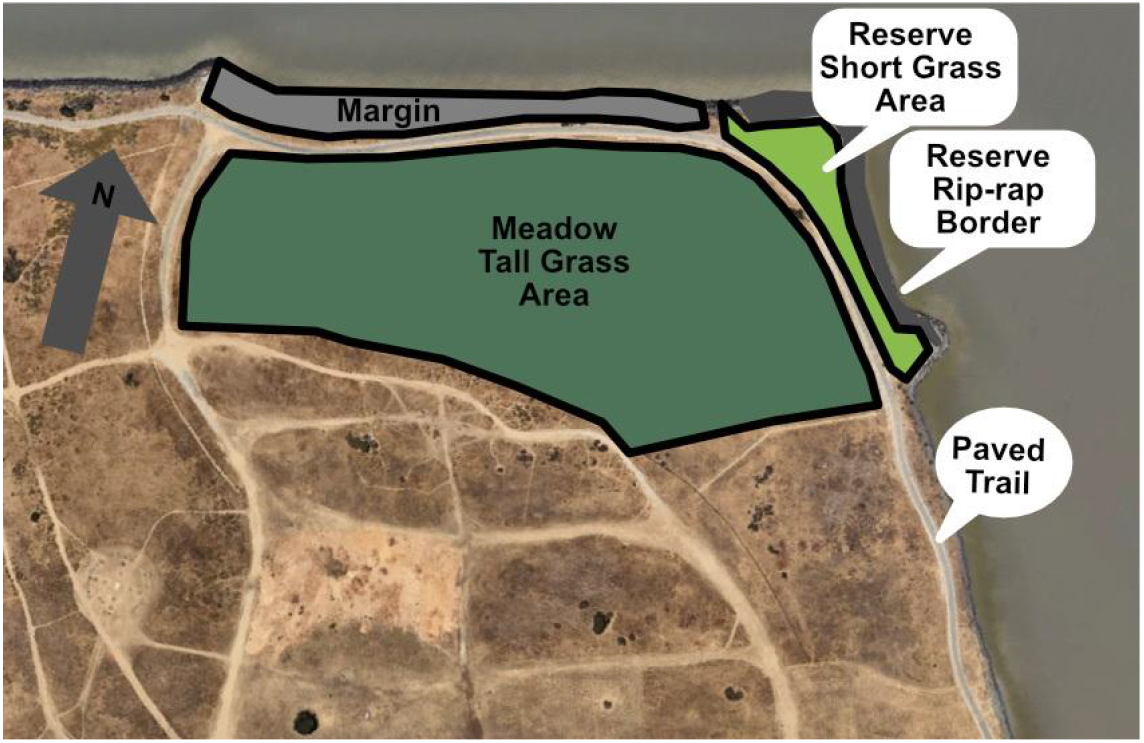

### (1) The Reserve

In the northeast corner between the paved trail and the water lies an area of about ¼ ha of flat grassland that since 2011 has been surrounded by a low fence. Known as the Burrowing Owl Reserve (or Sanctuary), this area contains two types of habitat.

(1a) A flat grassland with a scattering of fennel (*Foeniculum vulgare*) and coyote bushes (*Bacharis pilularis*) lies at the center of the Reserve. This portion is easily overseen from the paved perimeter trail and is usually mowed, in some years with removal of all taller vegetation. Owls present here settled as a rule in short grass or open gravel at spots located less than 1m from a ground squirrel burrow.

(1b) The north and east borders of the Reserve are rip-rap slopes. Owls settling on the east border may be completely or partially invisible from the path but can usually be seen with optics from a promontory about 100 m to the south. No owls were observed on the north border. No ground squirrel burrows are visible in the rip-rap. The spot where owls settled repeatedly in successive years featured a large dried California poppy bush growing in a crevice between stones. The plant formed a roof under which the birds took cover. When threatened, the owls dove into crevices and cavities in the rocks.

### (2) The Margin

In the margin between the paved trail on the north side and the water’s edge lies a sloping area of mixed vegetation. A screen of fennel borders the shoreline rip-rap. There is also a variety of ruderals and some coyote bushes. Vegetation here is never mowed. Owls were not observed on the soil portion of this area, but always on the rip-rap. Some could be seen easily from the path. Others settled behind a dense screen of fennel where it was a challenge to spot them.

### (3) The Meadow

To the south of the trail, opposite the Margin, lies a mostly flat grassland of about 3 ha that slopes up gently toward a ridge on its south side. This area is known in the park as “Protected Natural Area.” This grassland is never mowed in its interior. Vegetation here is a mix of native and introduced grasses and forbs with a handful of widely spaced coast live oak (*Quercus agrifolia*) shrubs less than 3m tall and coyote bushes less than 2m tall. There is a line of arroyo willows (*Salix lasiolepis*) on the south ridge. The height of the grasses and forbs here varies with the season and with rainfall, but typically exceeds 30 cm in winter. Many weeds reach or exceed 1 m. All owls observed in this area settled near ground squirrel burrows.

## RESULTS

### A. Owl numbers, arrival and departure dates, duration of stay

Based on the annual reports published by the docent program and later the daily reports on the chavezpark.org website, it is possible to reconstruct BUOW arrival and departure dates back to the winter of 2011-2012. These dates are set out in Table 1.

**Table 1:**
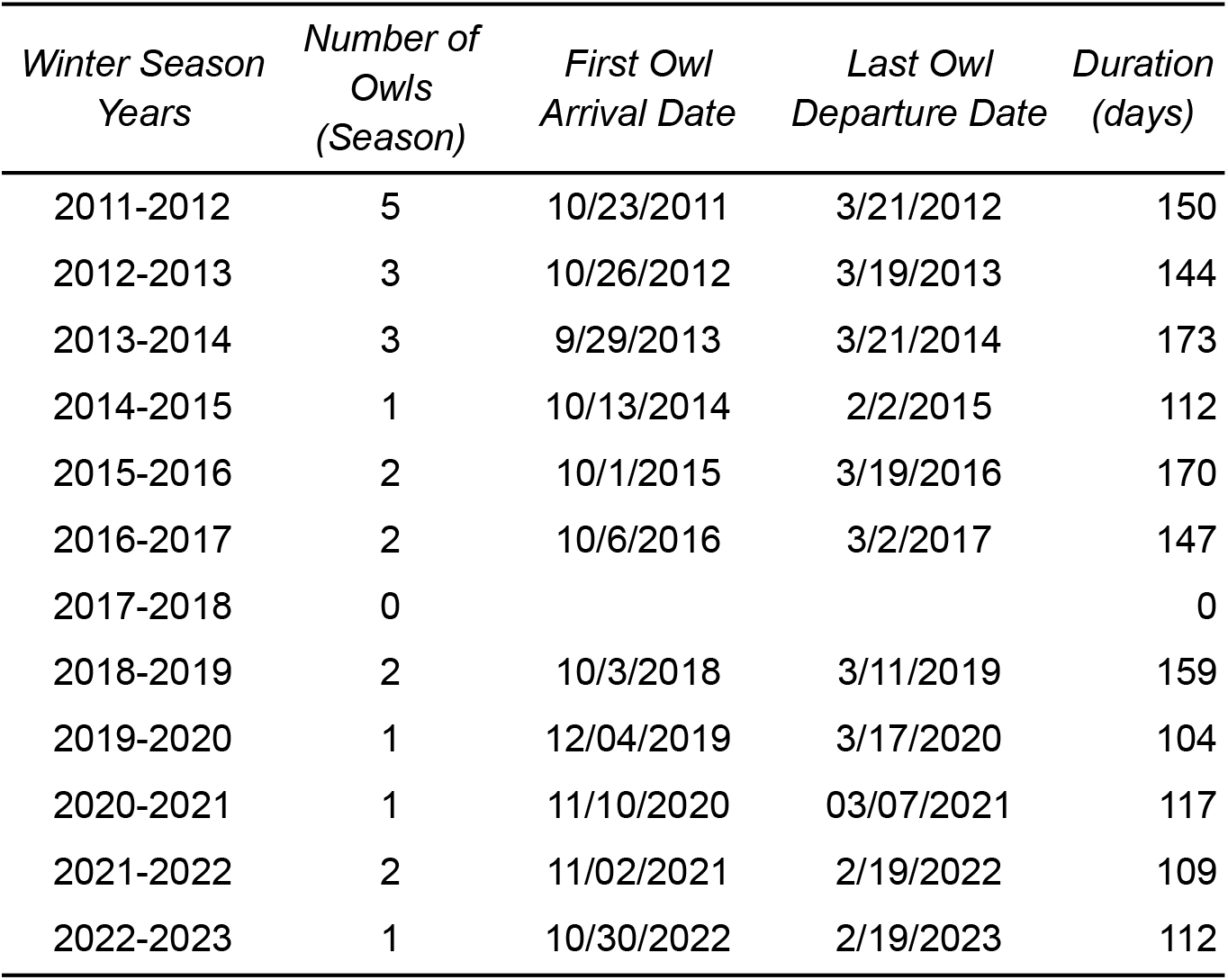
Burrowing Owls Settling for the Winter Season in Chavez Park, with Arrival and Departure Dates.

| *Winter Season*<br>*Years* | *Number of*<br>*Owls*<br>*(Season)* | *First Owl*<br>*Arrival Date* | *Last Owl*<br>*Departure Date* | *Duration*<br>*(days)* |
| --- | --- | --- | --- | --- |
| 2011-2012 | 5 | 10/23/2011 | 3/21/2012 | 150 |
| 2012-2013 | 3 | 10/26/2012 | 3/19/2013 | 144 |
| 2013-2014 | 3 | 9/29/2013 | 3/21/2014 | 173 |
| 2014-2015 | 1 | 10/13/2014 | 2/2/2015 | 112 |
| 2015-2016 | 2 | 10/1/2015 | 3/19/2016 | 170 |
| 2016-2017 | 2 | 10/6/2016 | 3/2/2017 | 147 |
| 2017-2018 | 0 |  |  | 0 |
| 2018-2019 | 2 | 10/3/2018 | 3/11/2019 | 159 |
| 2019-2020 | 1 | 12/04/2019 | 3/17/2020 | 104 |
| 2020-2021 | 1 | 11/10/2020 | 03/07/2021 | 117 |
| 2021-2022 | 2 | 11/02/2021 | 2/19/2022 | 109 |
| 2022-2023 | 1 | 10/30/2022 | 2/19/2023 | 112 |

These numbers and dates include only owls settling in for the season. Owls on stopover visits are included in Table 2 below. Two trends are apparent from Table 1.

**Table 2:**
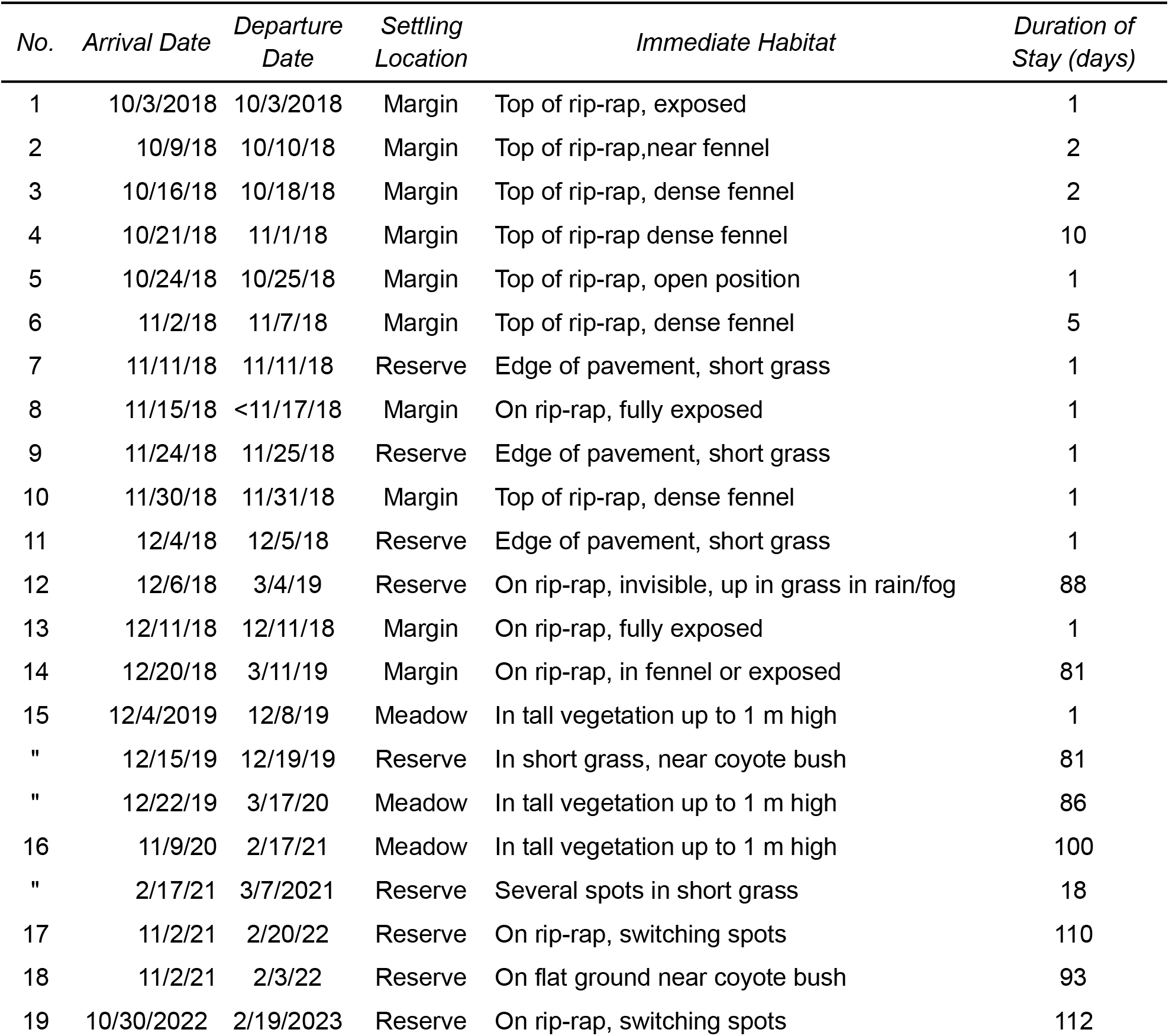
Burrowing Owls In Chavez Park During 2018-2023 Winter Seasons: Settling Location, Immediate Habitat, and Duration.

First, the number of Burrowing Owls spending the winter season in the park has declined. The reduction in this number is consistent with local and statewide trends. (Townsend and Lenihan 2003; Wilkerson and Siegel 2011; Center for Biological Diversity et al. 2024).

Second, the owls have tended to arrive later and leave earlier. The mean arrival date of the first owl in the first five years of observation (2011-2016) was October 11. In the last five years (2018-2023) the mean arrival date receded to November 3, a lag of 23 days.

Departure dates advanced, but much more narrowly. In the first five spring seasons, the mean last owl’s last day was March 9. In the last five seasons, the mean last day was March 3, an advance of six days.

The mean period when an owl was present during the first five years was 149 days. In the last five winters, the mean period of owl presence was 120 days, a reduction of almost a month.

### B. Choice of settling spots, micro-habitat

The Burrowing Owls seen in the park typically spent the daylight hours settled in one spot on the ground or other base, such as rip-rap. Accordingly,this study refers to settling or settlement spots, not perching or roosting. While some owls changed spots during the season, as noted below, the owls did not display the rapid daytime mobility typical of passerines.

The owls’ precise settling spots (immediate habitat) during the docent program period were not recorded in its annual reports. Precise settling spots, meaning the habitat in a diameter of about one meter around the bird, are available only for owls included in the Chavez Park Conservancy coverage. Almost all these observations are recorded as blog posts on chavezpark.org containing photographs and/or videos.

Table 2 shows the area that each visiting owl selected as its daytime settling spot, the immediate habitat at that spot, and the number of days the owl was observed in that spot.

The “Duration of Stay” numbers are subject to the reservation that the owls occasionally “took days off” when they could not be seen, but were later seen again in the same spot and are presumed to have remained present throughout.

Most owls remained in the same spot during the whole of their stay, but some owls moved. Owl No. 12 perched mainly in the rip-rap out of sight from the paved trail, but very occasionally came to the flat grassy area on foggy and rainy days. Owl No. 16 remained in the Meadow habitat but moved its location frequently, challenging its observers. In the last 18 days of its stay, Owl No. 16 relocated into the Reserve and moved among three different spots in that area. Owls No. 17 and No. 19 shuttled repeatedly between two different perches in the rip-rap, one where it could not be seen from the public path, and another where it was partly visible.

With these provisos, the table assembles habitat selections that owls made over a period of 797 days. Of this, 505 days (63%) were spent in the owl Reserve, 187 days (23%) in the Meadow, and 105 days (13%) in the Margin area.

The owl Reserve consists of two different habitats, the flat short-mowed grassland and the rip-rap shoreline. Of the 505 days the owls spent in the Reserve, 307 days (61%) were on the rip-rap, and 198 days (39%) were spent on the mowed grassland.

Looking at the owls’ presence as a whole, they chose to spend 198 of the 797 days (25%) on mowed grassland, 187 days (23%) in tall vegetation, and 412 days (52%) on rip-rap.

The owls’ habitat choices were split almost evenly between use of spots next to burrows and use of spots on rip-rap, where safety lay in cracks and cavities between stones. The owls used spots with burrows on 385 of their 797 days (48%) and spots other than with burrows 412 days (52%).

**Figure 3:**
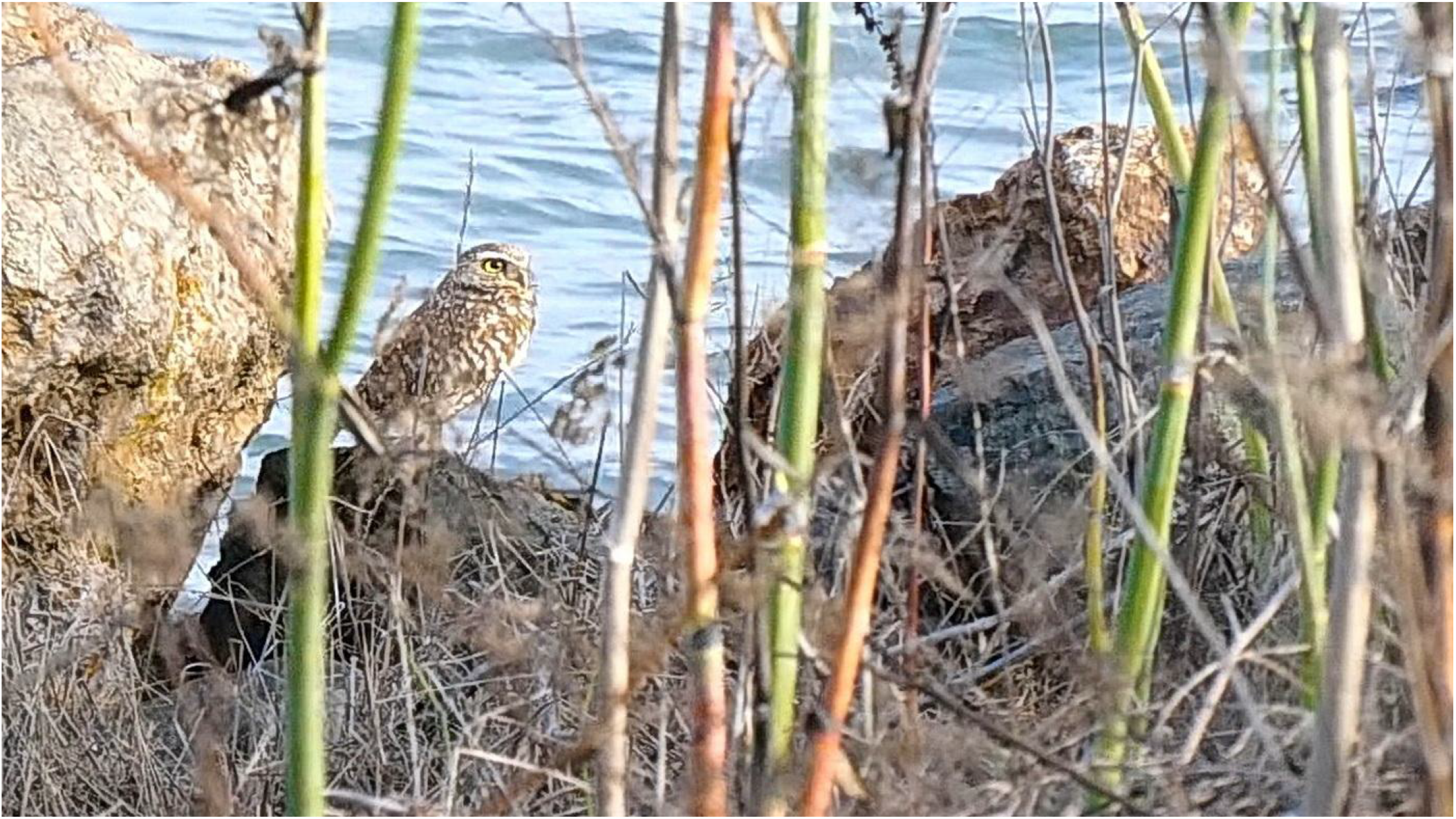
Owl No. 10 on rip-rap behind fennel screen in Margin area

**Figure 4:**
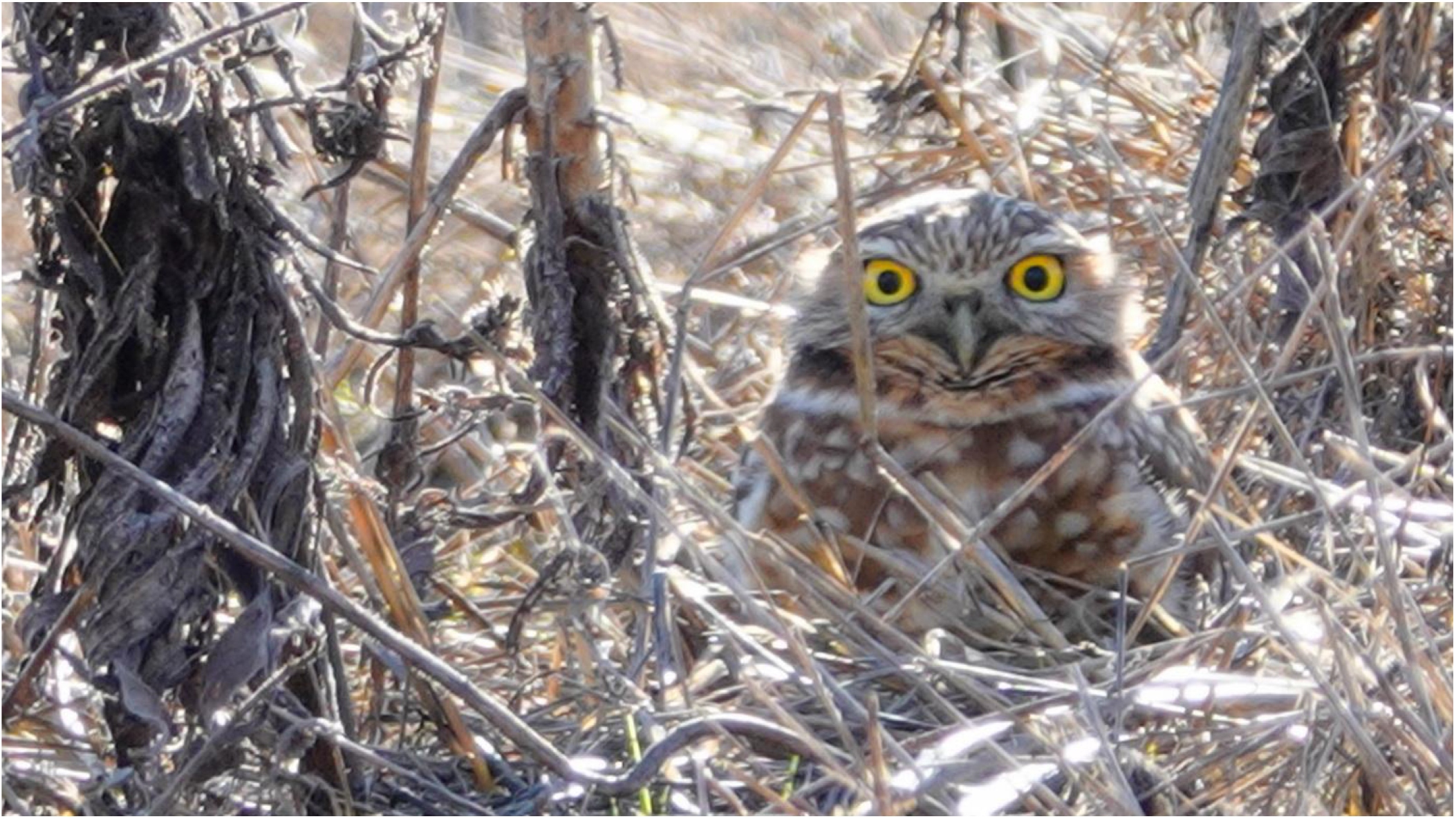
Owl No. 15 in tall vegetation in Meadow area

**Figure 5:**
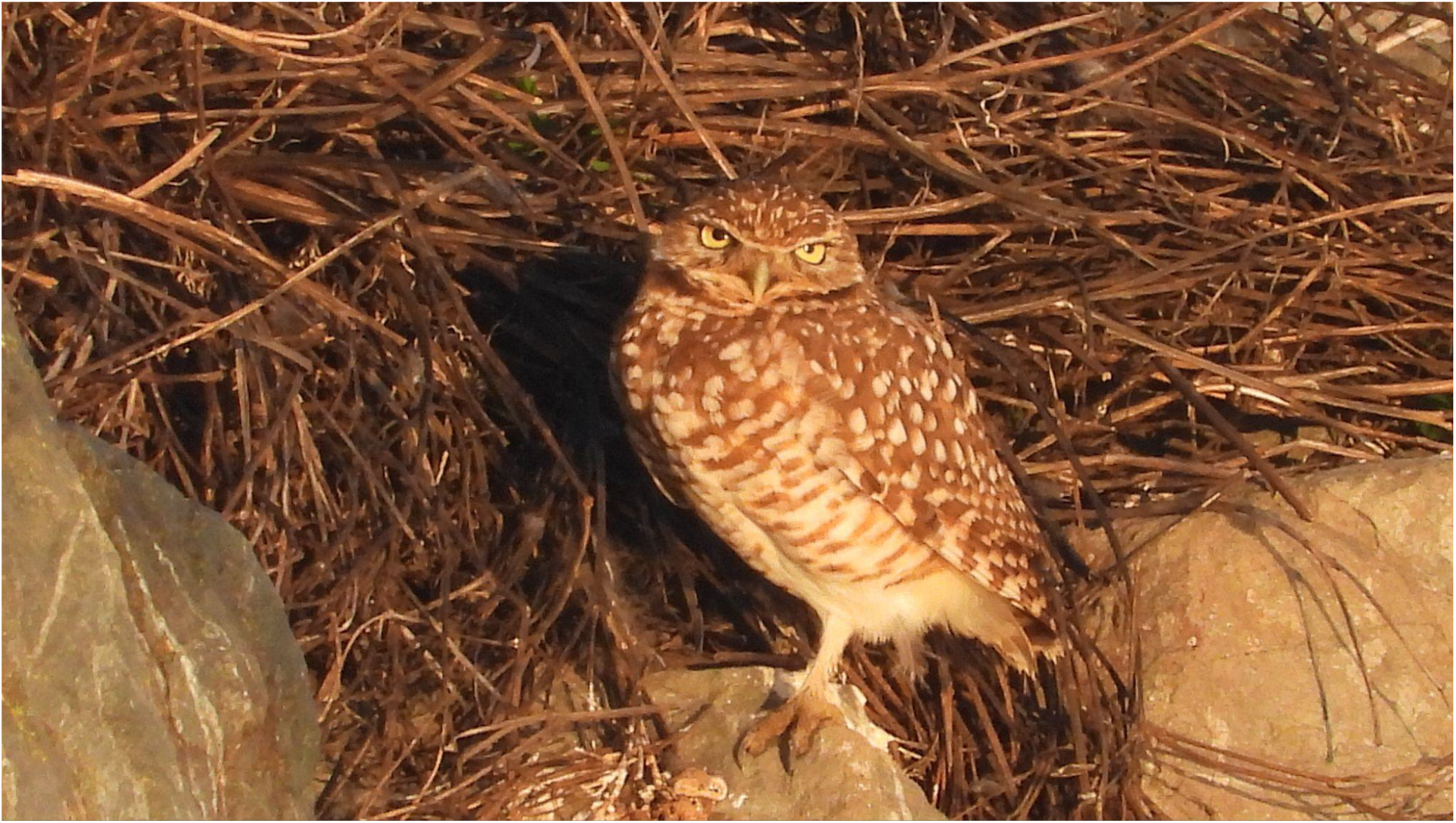
Owl No. 17 on rip-rap under dry shrub cover in Reserve

**Figure 6:**
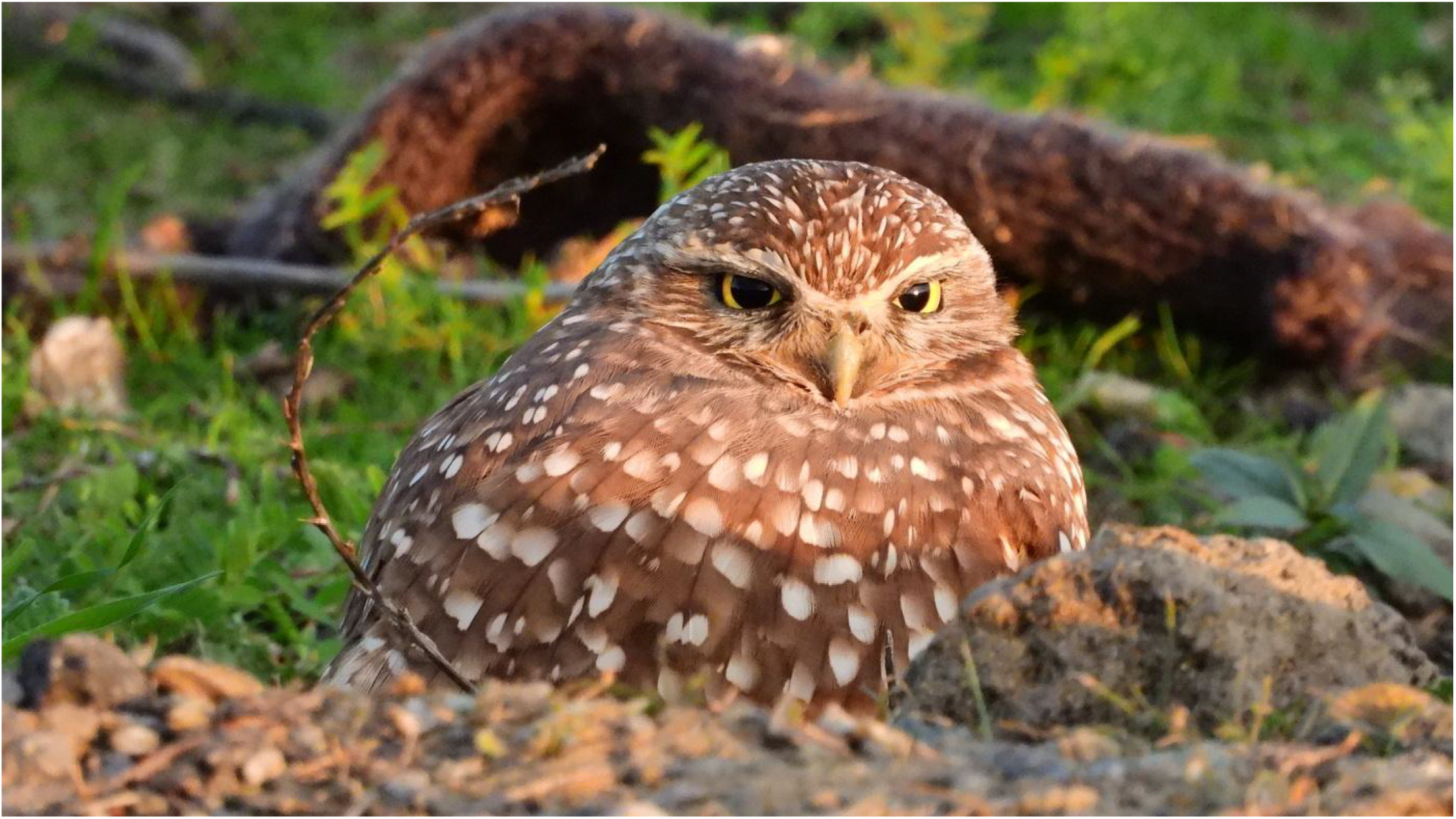
Owl No. 18 on short grass in Reserve

### C. Site Fidelity

There was little in the owls’ settlement patterns to suggest site fidelity. The strongest case was the choice of Owls No. 12 in 2018, No. 17 in 2021, and No. 19 in 2022 settling in the identical spot in the rip-rap on the east side of the Reserve -- the same stone under the same dried California poppy plant. It was not possible to determine whether they were the same individual.

### D. Behavior in Rain

During the winter seasons of 2021-2022 and 2022-2023, episodes of heavy rainfall termed “atmospheric rivers” occurred in the area, providing an opportunity to observe the birds’ behavior under intense and prolonged precipitation. These observations confirmed early reports that owls did not retreat into burrows during rain. (Butts 1973)

**Figure 7:**
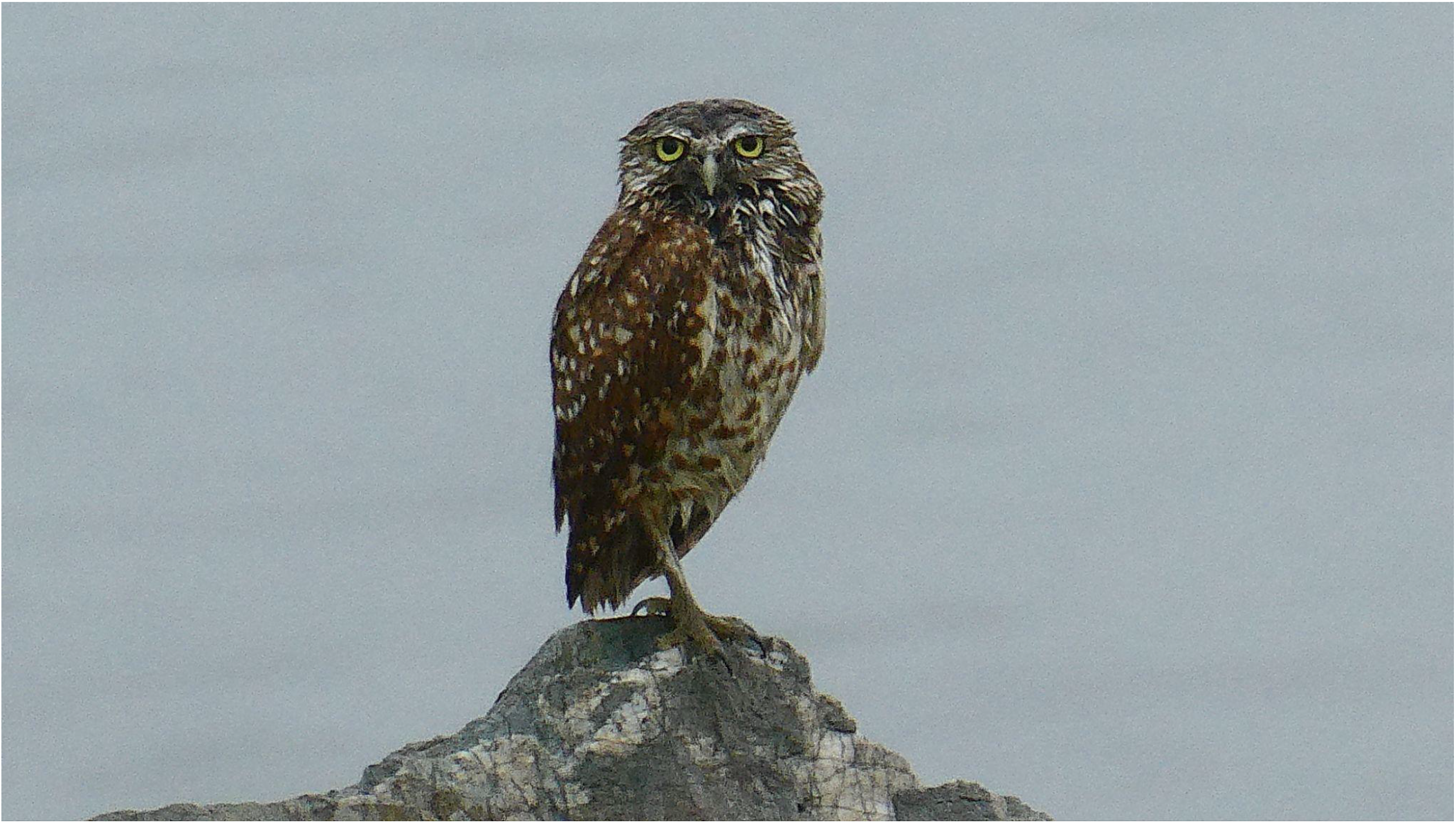
Owl No. 12 normally settled low in rip-rap, but perched on highest rock during heavy rainfall

The owls seemed to welcome exposure to rain. They sought out unsheltered spots and, in one case, perched on the highest, most exposed stone in the area during a downpour. Before long, the owls’ contour feathers were soaked. But the rain probably did not penetrate the dense, almost fur-like layer of very fine down next to their skin. From time to time, the birds shook themselves vigorously like wet dogs, sending raindrops scattering all around, and then continued bathing. But there was a limit. On the third day of rain, the birds were not to be seen. They emerged again when the sun returned. (Nicolaus 2019a)

**Figure 8:**
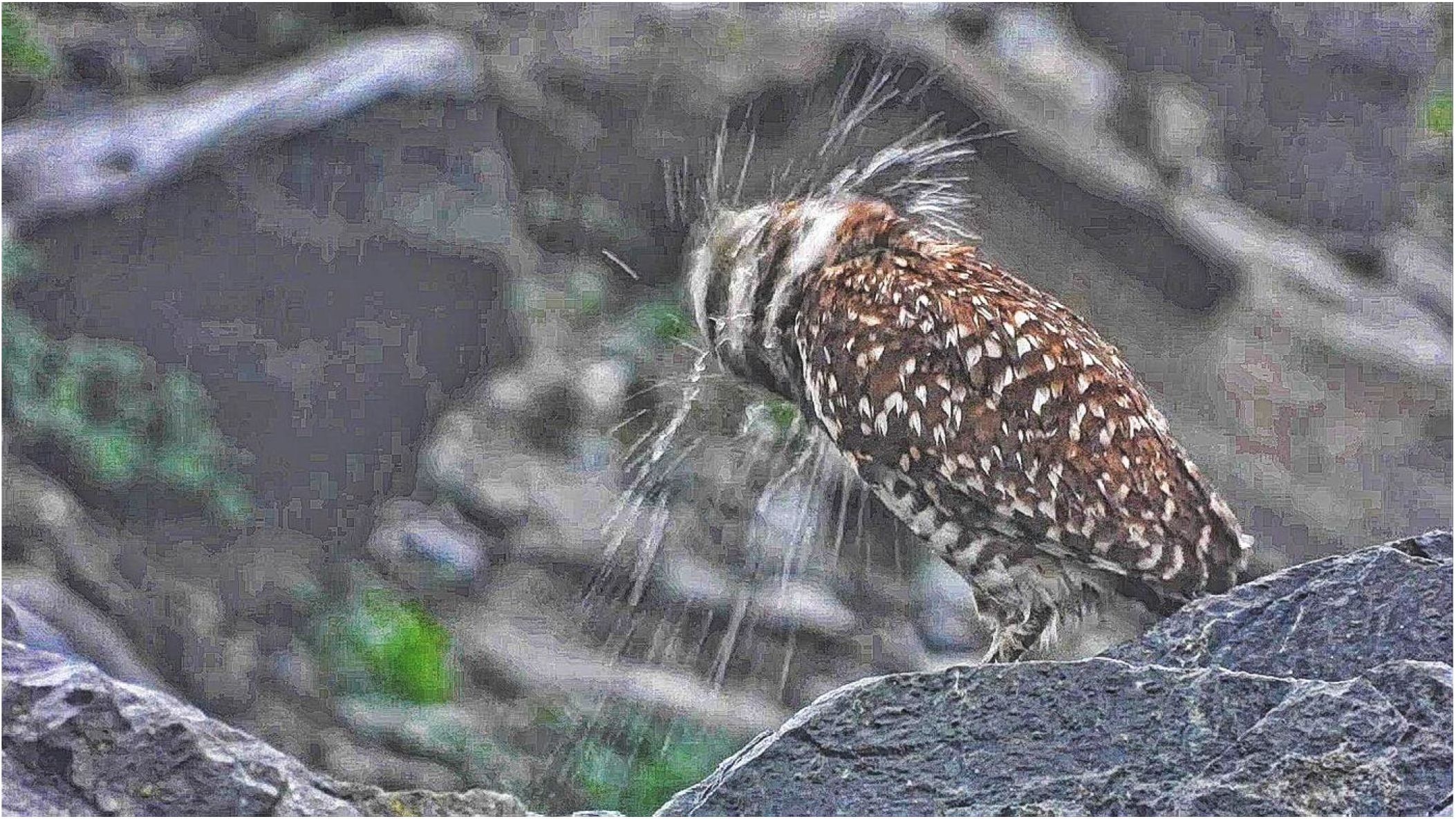
Owl No. 19 shakes off rain

In Fisher’s study, some owl chicks perished when nest burrows flooded. (Fisher et al. 2015) There is no indication that squirrel burrows flooded here. No burrow openings appeared filled with water. If burrows had flooded, the resident ground squirrels would be driven to the surface in large numbers, or they would drown and after the rain their numbers would be much reduced. But squirrels rarely appeared on the surface in the rain, and there was no noticeable shortage in their numbers afterward. Over the decades the rodents very probably engineered their burrows with adequate drainage. In any event, two of the owls here that chose exposure to precipitation had settled in the shoreline rip-rap, where cavities under the stones offered cover that could not flood, and where even the highest tides did not reach.

### E. Response to Human Exposure

I happened to be the second human being to encounter Owl No. 18 after its arrival in the park, and on my first approach this was a nervous bird. It didn’t flush and fly away when I set up my camera tripod about 7m away, but it looked at me with big eyes, stood on both feet ready to launch, and rotated its head frequently. By the next day it had calmed down considerably, and on the third morning my arrival and camera setup barely attracted its notice, as if saying, “Oh him again.”

A week after its arrival, this owl showed no fear of humans. At times, clusters of more than a dozen park visitors, including children and dogs, crowded the fence, talking loudly, gesturing, pointing, and taking photos. Occasionally a pickup truck belonging to Park management drove by or stopped on the path, with the driver viewing the owl. None of this led the owl to take evasive maneuvers or to display stress. Sometimes as the hubbub churned around it, the bird closed its eyes, yawned, or turned its back. This bird became the poster child of Burrowing Owls. It met the textbook description of owl habitat preferences: short grass or dirt, clear sight lines. Several hundred park visitors saw and photographed this owl. A brochure posted in a box near its spot dispensed 700 copies. People gave it names. Its fans featured it on every social media platform. I met visitors who came from more than 50 miles away to see it.

Although it arrived on the same day, Owl No. 17 chose a different habitat. It initially settled under a dry shrub in the rip-rap on the east border of the Reserve, where it was invisible from the perimeter trail. To see the bird one had to take up position on a promontory about 100m from the bird’s spot and use long-range optics. At that distance, observation did not distract the bird and it rarely glanced in the direction of the observer. But then after some weeks in its sheltered perch, the bird moved some 40m south to a new perch in the rip-rap, about 1m from a tall fennel bush. From there, at least its upper half and sometimes the whole bird could be seen from the perimeter trail, if an observer knew exactly where to stand. Crowds soon gathered at this spot to view the bird, which sat only about 10m from the trail.

The bird frequently looked in the direction of its noisy and agitated watchers but showed no sign of anxiety.

Human reaction to the owls was uniformly positive. An audio recording of humans viewing one of the owls displays a range of emotions from amazement to affection. (Nicolaus 2019b) The owls lived up to their reputation as charismatic birds. I know of no instance of a park visitor harassing an owl in person.

Some other owls, by contrast, displayed high levels of anxiety at the approach of a human. This was true particularly of some of the stopover owls seen during the winter of 2018-2019. Some of these displayed flight initiation distances of 15 or even 20m at the approach of a single walking human. There was no way to determine if these owls came from rural environments with scarce human presence, in contrast to more urbanized owls. (Carrete and Tella 2017) The conclusion, in any case, is that owls differ in their initial tolerance of the human presence, as in other qualities.

### F. Response to Predators

The broader park environment harbored a number of avian and mammalian predators active during the owls’ seasonal presence.

The Red-Tailed Hawk (*Buteo jamaicensis)* is the apex raptor in the park. It is the only demonstrated avian predator of ground squirrels in the park. I have observed two instances of a red-tail killing a squirrel. Both were messy scenes, with bloody bits of fur and bones strewn about. I have seen the red-tail fly low over the north side area where the owls settle, and I saw one red-tail perch on the rip-rap below one owl’s habitual spot, but I have never seen a red-tail attempt to attack an owl. The owls are 150 lean grams with sharp eyes and quick reflexes packaged in messy feathers. They settle among an active colony of bigger, fatter, slower earth-bound mammals weighing 280-700 grams. The dilution effect may well protect the owls from the apex raptor. (Bryan and Wunder 2014)

The Northern Harrier (*Circus hudsonius*) flies a low contour pattern that can easily surprise prey. Here, however, the birds generally have their backs to the water and the raptor will only approach over land, where it is more easily spotted. The harriers are infrequently seen in the park.

The White-tailed Kite (*Elanus leucurus*) seems to be a vole specialist and shows no interest in owls or squirrels.Owls sometimes watched a hovering kite with alertness, and at other times ignored it.

Barn Owls (*Tyto alba*) used nest boxes made available for them by the City in only one of the most recent five years. There were no reports of a Barn Owl attacking a Burrowing Owl here.

A Cooper’s Hawk (*Accipiter cooperii*) attacking a Burrowing Owl was captured on video. Owl No. 15 was settled in grasses in the Meadow. The hawk landed about 30cm away from the owl, peered at it curiously, and then stepped toward it in attack mode. The owl did a surprising thing. It charged the hawk with wings spread large and beak open wide, knocking the hawk on its heels. Then the owl turned 180 degrees and dove into the burrow at its feet. When the hawk recovered its composure, it took a few steps and flew off. (Nicolaus 2020b) The same owl also defied and drove off a group of 4 American Crows (*Corvus brachyrhynchos*) that were harassing it. The crows did not return.(Nicolaus 2020a)

In another encounter with a Cooper’s hawk, Owl No 19 dove for cover in the rip-rap moments before the hawk landed. The owl emerged again about 10 min later. (Nicolaus 2023)

The owls frequently scan the sky, tracking birds overhead, and seem to distinguish clearly between harmless birds (gulls, egrets, pelicans, etc.) and dangerous raptors. Where the owls see danger, they dive into their burrow or into cavities in the rip-rap in a fraction of a second. Two such occasions were captured on video. In each case, the owl remained out of sight for a matter of minutes and then resurfaced. (Nicolaus 2022)

In the memory of local observers, and in the records kept, there is no report of a Burrowing Owl in the park falling victim to an avian predator.

One or two abandoned cats (*Felis catu*s) inhabited and were fed in a woody grove about 600m away, but no cats were ever seen anywhere near the owl area. Raccoons (*Procyon lotor*), opossums (*Didelphis virginiana*), and skunks *(Mephitis mephitis*), seen only in very small numbers, likewise kept hundreds of meters away and did not frequent the north and northeast area of the park.

The apex predator in the park is pet dogs (*Canis lupus familiaris*). The Berkeley Municipal Code requires all dogs to be kept on leash except in posted areas. In 1998, the City designated a 7 ha area in the approximate middle of Cesar Chavez Park as an unfenced Off-Leash Area. The paved trail that passes by the Reserve, the Meadow, and the Margin areas is in an area where dogs must be on leash. Dogs are not permitted in the Meadow area at all, on leash or off. Nevertheless, some dog owners permit their dogs to run off-leash in all of these areas.

The first written report of Burrowing Owls wintering at Chavez Park dates from 2008 when an unlawfully off-leash dog charged an owl sitting in a burrow on the northeast corner and tried to dig it out. The owl disappeared. Volunteers belonging to the then Audubon Society chapter intervened and with the City’s cooperation erected a temporary plastic construction fence around the northeast corner area where owls had been settling. These fences are 1.2m (48 in) high. This was the beginning of the current Burrowing Owl Reserve. (GGAS 2012)

In 2010-2011 a wealthy donor financed a public art project in that area, consisting of two concrete seating fixtures, some low concrete retaining walls, and an Art Deco-style horizontal wire fence replacing the temporary fence. The permanent fence is lower, only 0.81m (32 in) high, composed of widely spaced parallel cables, with posts 1m (39 in) high. (Nicolaus, Martin 2018a) Dog incursions over or through the fence and over the low retaining walls were frequently reported during the docent program and frequently observed and photographed in more recent years, and have been recorded on video. (Nicolaus, Martin 2022a)

In late 2016, a burrowing owl was found dead in the park on a bench some distance from the owl Reserve. The cadaver was photographed but disappeared before a necropsy could be performed. The compressed but intact condition of the body ruled out an avian and pointed to a canine predator. (Nicolaus, Martin 2016)

In early February 2022, Owl No. 18, the “poster child” owl that had settled in the short grass easily visible from the path, abandoned its spot and hid behind a nearby stone. A video showed an injury to its left wing. The wingtip dragged on the ground and the bird walked with hesitation. Before anyone could organize an effort to save it, it disappeared. Several times in previous weeks, observers had seen off-leash dogs crossing the fence and flushing the owl. (Nicolaus, Martin 2022b)

The docent program asked docents to report the number of dogs that passed the Reserve on the paved trail, and whether they were leashed or not. The data show a rising trend in the number of off-leash dogs per hour, coinciding with a declining trend in the number of owls. (Table 3, Fig. 10)

**Table 3:** Docent Observations of Dogs and Owls in Cesar Chavez Park, 2011-2018.

| *Year* | *Total Dogs* | *On Leash Dogs Total* | *Off Leash Dogs Total* | *Percentage of Dogs Off Leash* | *Off Leash Dogs Per Hour* | *Owls* |
| --- | --- | --- | --- | --- | --- | --- |
| 2011-2012 | 776 | 646 | 130 | 17 | 1 | 5 |
| 2012-2013 | 619 | 464 | 155 | 25 | 1.5 | 3 |
| 2013-2014 | 898 | 723 | 175 | 19 | 1 | 3 |
| 2014-2015 | 401 | 197 | 204 | 51 | 1.2 | 1 |
| 2015-2016 | 219 | 177 | 42 | 19 | 0.7 | 2 |
| 2016-2017 | 306 | 195 | 111 | 36 | 2 | 2 |
| 2017-2018 | 476 | 151 | 325 | 68 | 4 | 0 |

**Figure 10:**
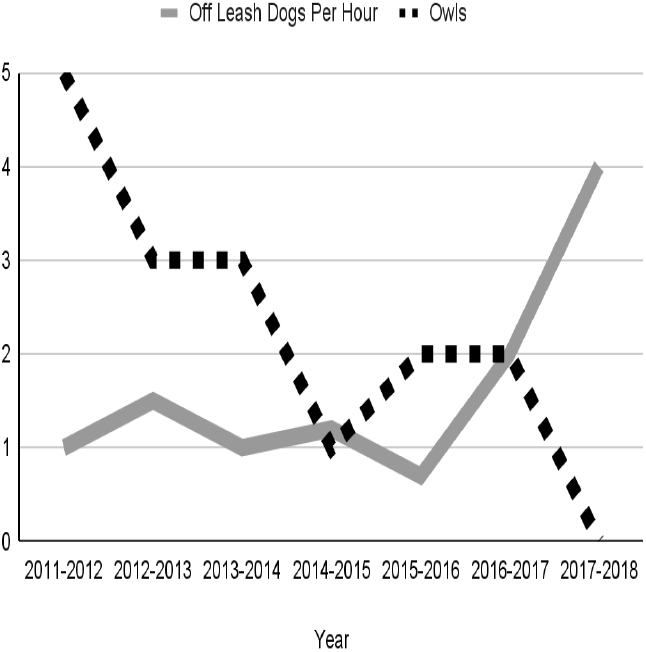
Off Leash Dogs Per Hour and Owls, 2011-2018

While these trends may well have lacked a causal relation, the rise in off-leash dog density did impact the number of volunteers participating in the docent program. “The lack of dog leash enforcement has led to several docents leaving the program,” according to the 2016-2017 annual report of the docent program. (GGAS 2017) The docent program tapered out after the winter of 2017-2018, when no owls were observed in Cesar Chavez Park.

Burrowing Owls watched but showed no alarm at dogs that could not reach them. In one instance recorded on video, an owl gave no reaction when a dog on leash passed, but went into high alarm mode, stretching full height, ready to fly, at the approach of a dog off leash. (Nicolaus, Martin 2018b) Numerous park visitors with dogs on leash stopped to observe visible owls, including Owl 18, without provoking signs of stress from the birds. One and sometimes two Burrowing Owls have wintered for several years on the edges of a major dog park, Pt. Isabel, a few km north of Cesar Chavez Park.

Effective fences stand between the owls and the dogs there, a distance sometimes of just a few meters.

### G. Owl Response to Predation Stress

Even when predation by loose canines does not result in death or injury to the owls, it stresses them. One ready and non-invasive measure of stress in owls is the frequency of their head rotations. The birds rotate their heads in order to check for danger. The greater their apprehension of danger, the more frequent their head RPM.

Two owls videotaped in February 2019 displayed sharply different head RPM. Owl No. 12 settled in the rip-rap of the owl Reserve, invisible to people and dogs on the perimeter path. This bird had never been disturbed by a dog while in residence in the park. The other owl, No. 14, settled in the Margin on the north side, where no barrier stood between it and the perimeter trail, about 10m away and 3m above its position. This owl sometimes hid in the fennel and at other times stood out on the rip-rap, easily visible.

Video recordings showed that this owl had been flushed by dogs at least three times during the preceding week.

Result: the owl that was never disturbed by a dog clocked 8 head rotations per minute. The owl that had been repeatedly disturbed turned its head 46 times per minute. Carried on throughout the daylight hours that would mean 2,760 rotations per hour or more than 25,000 rotations per day. This level of muscular effort has a caloric price that likely rendered the bird less fit than optimum for its spring migration and reproduction. (Nicolaus 2019c)

## DISCUSSION

Given the slim wintering literature, the Burrowing Owls in Cesar Chavez Park on the Berkeley waterfront may fill in some gaps. Certain aspects dovetail with known trends, notably the decline in their numbers over the past two decades, and very likely their later fall arrival and earlier spring return, possibly as an adaptation to climate warming in northern breeding habitats. (Barr et al. 2023; Sales and Parrott 2024)

A case of possible site fidelity was seen here, with three instances of an owl or owls settling in the identical spot in rip-rap. But the majority of the owls showed no consistent settling spot pattern, other than that all lay on the north side of the park.

The most notable finding is the owls’ choice of micro-habitat for settling spots. It is a common theme in the literature that Burrowing Owls prefer to nest in short or absent vegetation with clear sight lines. The classic statement, oft-quoted, is probably that of Elizabeth Haug’s MS thesis: “All agree the major criteria essential for nesting habitat are openness of site, availability of nest burrows and short grass.” (Haug, Elizabeth A. 1985) This observation of breeding ecology has been enlarged into a general prescription for habitat management in the winter season. In Cesar Chavez Park, a permanent sign next to the Reserve says that “They favor areas where low, sparse vegetation allows them to see predators from a distance… “ Following this formula, park management routinely short-mowed the grass and removed all taller vegetation in the Reserve area ahead of the owls’ expected arrival date, so that the area resembled a moonscape. Yet the visiting owls here overwhelmingly favored areas of tall vegetation and rocky areas where they were difficult to see, selecting short grass only in a minority of cases.

Do these results mean that the owls’ wintering habitat choices are radically different from their preferences in breeding season? This may be true to some extent, but a close review of the literature indicates that their actual nest site habitat selections were far more diverse than the “short” and “open” formula indicates.

Even in Haug’s own study, there were exceptions. Two owl pairs nested in dense grass about 75 cm high, and a third pair nested in a spot surrounded by tall sheds and bleachers. (Haug, Elizabeth A. 1985)

In a South Dakota nest setting, while vegetation was short at the start of the breeding season, “annual forbs eventually dominate the burrow” and “provide more concealment for emerging young owls.” (MacCracken, Uresk, and Hansen 1985)

In south-central Idaho, occupied burrows had a greater vegetation cover than others. (Rich 1986)

In a Colorado military reservation, Burrowing Owl vegetation preference varied by year. (Plumpton 1992)

In an Alberta study, there was no difference in habitat between prairie dog burrows occupied by owls and those not occupied. The researcher suggests that observation of low vegetation density “may reflect the choices made by burrowing mammals more than burrowing owls.” Prairie dogs (*Cynomis spp*.) are aggressive herbivores that clear their surroundings of vegetation. (Schmutz 1997)

In a park setting in Northern California, 32 percent of nest burrows were in tall grass, and at a nearby airfield, 15 percent were in tall grass. (Trulio 1997)

In Saskatchewan crop lands, “The owls’ nests were not selectively placed in short vegetation … Here owls also nested in the non-agriculturally used strips of medium to tall grass between fields and other features such as ponds, granaries and roads.” (Clayton and Schmutz 1999)

A Wyoming study found variability among landscape and habitat use, concluding that Burrowing Owls do not have extremely specific habitat requirements. (Lantz, Smith, and Keinath 2004)

Fledgling mortality was especially high in nesting areas with sparse cover and “lower in regions with an abundance of escape cover.” (Morgan Davies and Restani 2006)

In a study on a military base in California, researchers found owls nesting in tall grass areas, and hand-mowed the grass around the burrows to facilitate observation. (Fisher, Joshua B. et al. 2007) Here it was the observers, not the observed, that preferred open view sites.

In a desert setting where tortoises and kit foxes rather than prairie dogs created burrows, the owls preferred sites with a greater cover of creosote bush surrounding the nest site, where they were difficult to detect. (Crowe and Longshore 2013)

In Canadian prairie lands generally, there was no consistent preference for open habitat. “The one commonality of burrowing owl nest- and foraging-habitat selection is that the owls appear to be ambiguous to or possibly prefer heterogeneity of vegetation types.” The researcher concluded that “the Burrowing Owl is quite flexible in its habitat requirements and can make a living almost anywhere within their range.” (Scobie 2015)

In eastern Kansas, vegetation surrounding burrows stood at about 10 cm in May but exceeded 40 cm in mid-July when chicks fledged. (Herse 2016)

In a wind turbine area of Northern California, a researcher visited Burrowing Owl nests located in waist-high grass and others in rapeseed (*Brassica napus*) over his head. “The idea that Burrowing Owls don’t nest in tall grass is a myth. We think they don’t breed there because we can’t see them. But if we know where to look we can find them there.” (Shawn Smallwood, personal communication, 2023).

The Western Burrowing Owl is dependent on others to dig burrows for it, and this may restrict its choice of habitats. Burrowing Owl species that dig their own burrows, like *Athene cunicularia floridana*, can nest wherever they like. (Millsap and Bear 2000) In Florida, two pairs of owls successfully fledged young in large patches of tall vegetation. (Mrykalo 2005) In the Argentine pampas, where the Burrowing Owl species also dig their own burrows, the owls “showed little selectivity of nest sites and nest patches, thus reinforcing the idea that they are habitat generalists.” (Martinez et al. 2017)

Recent studies of wintering owls also show the owls selecting micro-habitats other than short grass with clear sight lines.

Wintering owls in the Great Plains used culverts, junk piles, or hid in dense vegetation. (Clayton and Schmutz 1999)

In southern Texas, 74 percent of observed owls roosted in manufactured culverts of steel or stone, where it was difficult to see them. (Williford et al. 2007; Woodin, Skoruppa, and Hickman 2007)

A radiometry survey of owls migrating from Canada and wintering in Texas and Mexico found that “land-cover types around roosts were highly variable and less open than breeding habitat in Canada.” One roost was in a mammal burrow, one in a woodpile, and five were surrounded by thick shrubland or similar vegetation providing dense cover. All habitat plots included trees and shrubs. (Holroyd, Trefry, and Duxbury 2010)

In the northern Sacramento Valley of California, in fall and winter, about a third of natural burrows were located in grasses or forbs more than 16 cm high. Around active artificial burrows, 61 percent were set in grasses or forbs 22 cm high or higher, enough to completely conceal a standing owl. (Ocken 2018)

A study of owls wintering in central Mexico showed that roosting selections were highly variable, ranging from anthropogenic cavities to trees. The authors concluded that “Burrowing Owls have flexible requirements for roost sites in winter.” (Valdez-Gómez et al. 2018)

The owls wintering In Cesar Chavez Park chose to settle on the mowed grassland only 25% of the time. And nearly half of that (12% of the total) was accounted for by one owl, No. 18, which spent all 93 days of its presence in that habitat. As noted, this bird was readily visible from the perimeter trail and became the star of a fan club with hundreds of human admirers. But among wintering owls in Cesar Chavez Park, this choice was an outlier. And there were consequences. The area where it settled had inadequate protection against loose canines, and the bird’s stay was cut short by a disabling injury.

There are two sides to clear sight lines: the owl can see predators, but predators can see the owl. Like vast numbers of other prey species, the owls can benefit from vegetative cover. (MacCracken, Uresk, and Hansen 1985)

## CONCLUSIONS

Unlike remote wilderness, the urban park habitat is an intensely human-managed biome. Management cannot control the weather, but it controls the physical base, the soil and stones, the selection and structure of vegetation, and the infrastructure of human access. When this habitat is visited by avians that are of special concern, management decisions need to be based on a sound understanding of the ecology that ensures the species’ survival.

The extension of concepts derived from a species’ breeding habitat to its wintering habitat is normally a fraught operation. All the more is this true when the analysis of breeding habitat is based on one selected aspect, leaving out others. So, while it is clearly true that Burrowing Owls often select short-grass, high-visibility micro-habitats for nesting, the evidence is quite strong that they can and do also select breeding habitats in tall grass and other spots where humans cannot easily see them. The owls are habitat generalists, able to breed and raise young in a great diversity of ecological sites.

In the case of Burrowing Owls, there is a temptation to project human preferences onto the birds. Since we prefer owls in spots where we can easily see them, we assume that owls also prefer such spots. In the words of Shawn Smallwood, who has studied the owls for decades, “We think they don’t breed there [in tall grass] because we can’t see them. But if we know where to look we can find them there.”

We may even deliberately cut the grass short around their nest burrows, as researchers did in a California military base (Fisher, Joshua B. et al. 2007), as if the birds had made a mistake nesting in tall grass and we were correcting them and doing them a favor.

To reinforce the human projection, we say that the owls’ choice of high-visibility habitat is motivated by their desire to see approaching predators. This might be plausible if the owls were big apex raptors not needing to hide. In reality, the owls are very small raptors. They figure more as prey in avian ecology than as predators. Normally, prey species value habitat that offers shelter and concealment. The thesis that the owls favor high-visibility sites, when translated into ecological management choices, exposes the owls to danger and limits their defenses.

Ecological management based on unsound estimates of species requirements can turn a habitat into an ecological sink -- a place where conditions promote their extirpation. For the conservation of the wintering owl population in this park habitat, two recommendations seem most urgent. First, the seasonal Burrowing Owl preserve cannot continue to operate without installation of an effective dog barrier. Even if a bird is not killed, assaults by canine predators cause stress that diminishes their migratory and reproductive potential.

Second, the eradication of all but the shortest vegetation in an effort to create an “open site” with clear sightlines does not serve the birds. It may serve human observers to see the owls if they come there, but the exceptional owls that choose this setting are exposed to unreasonable predatory pressure. The park is not a zoo dedicated to animal display. The park needs to serve as an ecological niche also for owls that prefer a more sheltered and secure micro-habitat. If that makes it harder for us to see them, so be it.

## ACKNOWLEDGEMENTS

This study owes much to the contributions of park visitors who spotted, photographed, or reported Burrowing Owls, including Andre Bourgoyne, Jutta Burger, Juan Camacho, Della Dash, John Davis, Anthony DeCicco, Ian Flaherty, Sarah Flaherty, Donna Hom, Bob Huttar, Sheila Jordan, Anna Klafter, Louis Kruk, James Kusz, Laura Lange, Mary Law, Steve Lefkovits, Rick Lewis, Mary Malec, Alex Milhano, Linda Morris, Johanna Naederhouser, Keenan Quan, Christopher R., Phil Rowntree, Rachel S., Shawna S., Sara Sato, Jacob Several, Matt Shogren, Brendan Tidd, Allison Tom, Noreen Weeden, Shiyang Wu, Suzanne York, Tianxi Zheng, and Sam Zuckerman.

